# Soluble Immune Factor Profiles in Blood and CSF Associated with LRRK2 Mutations and Parkinson’s Disease

**DOI:** 10.1101/2025.03.20.644460

**Authors:** Roshni Jaffery, Yuhang Zhao, Sarfraz Ahmed, Jackson G. Schumacher, Jae Ahn, Leilei Shi, Yujia Wang, Yukun Tan, Ken Chen, Hussein Tawbi, Jian Wang, Michael A. Schwarzschild, Weiyi Peng, Xiqun Chen

**Affiliations:** Department of Biology and Biochemistry, University of Houston, Houston, TX 77204; Aligning Science Across Parkinson’s (ASAP) Collaborative Research Network, Chevy Chase, MD 20815; Department of Neurology, Mass General Institute for Neurodegenerative Disease, Massachusetts General Hospital, Harvard Medical School, Boston, MA 02114; Department of Melanoma Medical Oncology, The University of Texas MD Anderson Cancer Center, Houston, TX 77030, USA; Department of Bioinformatics and Computational Biology, The University of Texas MD Anderson Cancer Center, Houston, TX, USA; Department of Biostatistics, The University of Texas MD Anderson Cancer Center, Houston, TX, USA

**Author notes:** Contributed equally (Roshni Jaffery, Yuhang Zhao).

**Keywords:** *LRRK2*, Parkinson’s disease, immune regulators, soluble immune biomarkers, SDF-1 alpha, TNF-RII

## Abstract

**Background and Objectives:** Mutations in the Leucine-rich repeat kinase 2 (*LRRK2*) gene are one of the most common genetic causes of Parkinson’s disease (PD) and are linked to immune dysregulation in both the central nervous system and periphery. However, peripheral and central profiles of soluble immune factors associated with *LRRK2* mutations and PD have not been comprehensively characterized. Using serum and CSF samples from the LRRK2 Cohort Consortium (LCC), this study aimed to probe a broad range of soluble immune biomarkers associated with *LRRK2* mutations and PD.

**Methods:** We investigated the levels of soluble immune regulators in the serum (n=651) and cerebrospinal fluid (CSF, n=129) of *LRRK2* mutation carriers and non-carriers, both with and without PD. A total of 65 cytokines, chemokines, growth factors, and soluble receptors were assessed by Luminex immunoassay. A multivariable robust linear model was used to determine levels associated with *LRRK2* mutations and PD status, adjusting for age, sex, and sample cohort. Correlations were assessed using the Spearman correlation coefficient. *LRRK2* G2019S knock-in mice were used to validate the associations identified in the LCC.

**Results:** In this extensive discovery cohort, we identified several elevated serum immune regulatory factors associated with *LRRK2* mutations. In particular, serum stromal cell-derived factor-1 alpha (SDF-1 alpha) levels, as supported by findings in LRRK2 G2019S knock-in mice, and tumor necrosis factor receptor II (TNF-RII) were significantly increased after multiple comparison adjustment. In contrast, *LRRK2* mutations were associated with reduced soluble immune markers, including BAFF, CD40-Ligand, I-TAC, MIP-3 alpha, NGF beta, and IL-27 in CSF. Those with clinically diagnosed PD, with or without *LRRK2* mutations, did not show strong signals in serum but reduced inflammatory analytes in CSF, including MIF, MMP-1, CD30, Tweak, and SDF-1 alpha. In addition, we found that the serum levels of these soluble immune factors display varied correlations with their corresponding CSF levels.

**Discussion:** This study highlights distinct immune profiles associated with LRRK2 mutations and PD in the periphery and CNS. Serum levels of SDF-1alpha and TNF-RII were elevated in LRRK2 mutation carriers, while CSF immune markers were reduced. In PD, irrespective of LRRK2 status, reduced CSF inflammatory analytes and weak serum signals were observed. These results provide insight into immune dysregulation linked to LRRK2 mutations. If replicable in independent datasets, they offer potential avenues for biomarker and therapeutic exploration.

## Introduction

Parkinson’s disease (PD) is a common age-related neurodegenerative disorder manifested clinically by resting tremor, rigidity, bradykinesia, and gait instability, as well as non-motor features ranging from hyposmia to depression. Although the exact etiology remains unclear, evidence suggests that a combination of genetic and environmental factors contributes to its development.^1,2^ Among the known genetic risk factors, the leucine-rich repeat kinase 2 (*LRRK2*) gene has been identified as one of the genetic factors associated with PD. Mutations in *LRRK2* are associated with an increased risk of both familial and sporadic forms of PD. *LRRK2* PD is associated with variable clinical presentations that are often indistinguishable from idiopathic PD. A few studies reported slower disease progression rates of *LRRK2* PD.^3,4^ LRRK2 is a complex and multifunctional enzyme and is involved in a variety of cellular processes, such as cytoskeletal dynamics, vesicle trafficking, and autophagy, making it a critical player in maintaining cellular homeostasis and function. Mutations of *LRRK2*, including the most prominent G2019S, often lead to increased kinase activity, which is thought to contribute to the degeneration of dopaminergic neurons, a hallmark of PD, though the exact mechanisms by which mutations in *LRRK2* contribute to PD are still being elucidated.^5^

Growing evidence suggests that immune dysregulation may play a crucial role in the pathophysiology of PD. Neuroinflammation, characterized by the activation of microglia and astrocytes, as well as infiltration of peripheral immune cells into the central nervous system (CNS), has been implicated in the progressive neurodegeneration observed in PD.^6,7^ Proinflammatory cytokines and chemokines released by activated immune cells contribute to the degenerative cascade, exacerbating neuronal damage and promoting disease progression.^6–7^ Indeed, alterations in cytokine and chemokine profiles have been reported in PD. Elevated levels of cytokines such as interleukin (IL)-1beta, IL-6, IL-18, and tumor necrosis factor (TNF)-alpha have been detected in the CSF of PD patients compared to controls.^8–10^ Additionally, increased levels of cytokines such as TNF-alpha, IL-1 beta, IL-6, and IL-10 in the serum or plasma of PD patients have also been reported, suggesting systemic immune dysregulation in PD.^11–13^

LRRK2 is frequently expressed in many immune cells in the CNS and the periphery, including microglia, monocytes, T cells, and B cells.^14^ Beyond PD, variations in *LRRK2* are also linked to an increased risk of inflammatory bowel disease and greater susceptibility to bacterial infections, highlighting the role of LRRK2 in regulating immune and inflammatory responses.^15^ Elevated levels of serum IL-1 beta, TNF-alpha, IL-6, IL-10, and MCP1 have been observed in asymptomatic *LRRK2* G2019S carriers.^16^ In PD, increased LRRK2 expression was observed in innate and adaptive immune cells.^17^ Studies have shown a positive correlation between LRRK2 expression and IFN-gamma, TNF, and IL-2 expression in T cells from PD patients, suggesting that LRRK2 may further exacerbate immune dysregulation in PD.^17^ Inflammatory responses driven by increased *LRRK2* kinase activity is proposed to play an early role in PD pathophysiology.^18^ Other studies suggest that inflammation in *LRRK2* PD could be further intensified by relative impairment of anti-inflammatory cytokines like IL-10.^19,20^ A comprehensive overview of immune dysregulation and cytokine alterations in PD and *LRRK2* carriers can be found in recent review articles.^8,13,14^

To further characterize peripheral and central immune dysregulation associated with *LRRK2* mutations and PD, we determined levels of 65 cytokines, chemokines, growth factor targets, and soluble receptors/ligands, which have reported roles in regulating immune and inflammatory responses in serum and CSF samples from participants enrolled in the LRRK2 Cohort Consortium (LCC). We identified and compared the shifts and trends of the analyte concentrations found in the serum and CSF of subjects with mutations in *LRRK2* without PD, with idiopathic PD lacking a *LRRK2* mutation, with *LRRK2* PD, and without the *LRRK2* gene mutation and PD. Our findings in this discovery cohort and our validation using *LRRK2* G2019S knock-in (KI) mice provide insights into potential immune-related biomarkers in serum and CSF associated with PD and *LRRK2* mutation status.

## Methods and Materials

### Summary of reported literature of cytokine study using Generative Pre-Trained Transformer (GPT) models

To analyze the associations between cytokine concentrations and PD, including those carrying mutations of *LRRK2*, we used a generative pre-trained transformer (GPT) model to extract cytokine-related information from the literature.^21^ GPT models are a type of large language model (LLM) that learns patterns and structures from existing data, such as text and images, using deep learning and applying them to analyze and generate content.^22^ Leveraging OpenAI’s 4o GPT model, we extracted cytokine information through the following steps: The most relevant papers were retrieved from the PubMed Central (PMC) database through an application programming interface (API) using keywords: (cytokine name) AND (LRRK2 AND Parkinson) OR (LRRK2) OR (Parkinson). The search was limited to a maximum of 50 papers per cytokine. Publication years were retrieved through PubMed API. Full texts, excluding references, were processed by the subsequent GPT model.

### GPT Prompt engineering and postprocessing

We structured the instructions for the GPT model into two categories: custom prompts and system prompts. In custom prompts, instructions were given to confirm the presence of the cytokine of interest, define disease context, host type, measurement site, association with *LRRK2*, mutation variants, association with PD, and, most importantly, supportive evidence directly extracted from the literature. The system prompt ensured the quality and consistency of model response by providing definitions of *LRRK2* mutation and association and a list of alternative names for cytokines. Scores of 1 were given to positive associations between higher cytokine concentrations and PD or the presence of *LRRK2* mutation, −1 to negative associations, and 0 to no associations. The associations were inferred by the GPT model based on texts in the literature. The model was instructed to infer a positive association if 1) a cytokine’s concentration is higher in PD or PD model hosts if provided, or 2) in the presence of the cytokine, the odds ratio between PD and non-PD is greater than 1 and the lower bound of confidence interval of greater than 1; or) the inhibition of the cytokine is associated with increased risks of PD. We included quality control steps ^23^ in custom prompts to verify the extracted content against the literature and make corrections where necessary. The model’s performance was improved through iterative validation of the extracted responses, with prompt modifications applied based on the validation results. Each iteration of manual evaluation of model accuracy involved randomly selecting 10 to 30 PubMed IDs (PMIDs) and validating model responses of cytokine extracted, disease type, support for LRRK2 mutation, support for association with PD, host, and sites. Entries with no cytokine information found were removed from the analysis. For association analysis, two entries with disease types equal to traumatic brain injury (TBI) and depression in PD (dPD) were removed due to irrelevance.

### Human Samples

This project utilized samples from the LCC provided by the Michael J. Fox Foundation (MJFF). LCC is a large dataset established in 2009 with coordination and funding from the Michael J. Fox Foundation for Parkinson’s Research (MJFF). The LCC comprises individuals diagnosed with idiopathic PD (*LRRK2*−/PD+), unaffected noncarrier controls (*LRRK2*−/UC), carriers of pathogenic *LRRK2* mutations who also have PD (*LRRK2*+/PD+), and *LRRK2* mutation carriers without PD symptoms (*LRRK2*+/UC). Additional details about the LCC have been published and can be accessed at michaeljfox.org/data-sets and michaeljfox.org/lccinvestigators.

We received a total of 651 serum samples and 129 CSF samples in 4 sample cohorts from the LCC. Samples were stored at −80°C until analyzed. This study uses deidentified samples and clinical data from the LCC. Human subjects research is exempt as defined by Title 45 Code of Regulations (CFR)46.

### Bioplex Multiplex Immunoassay

The levels of 65 cytokines, chemokines, growth factor targets, and soluble receptors in serum and CSF samples were determined by the ProcartaPlex human immune monitoring kit (EPX650-10065-901, ThermoFisher Scientific, Waltham, MA) on a Luminex-200 system according to the manufacturer’s instructions. The samples were processed in prearranged batches, each containing approximately balanced numbers of samples from different groups. A 25-microliter volume of each serum or CSF sample was utilized.

The 65 pre-determined cytokines, chemokines, and growth factor targets, and soluble receptors include: G-CSF (granulocyte colony-stimulating factor, also known as colony-stimulating factor 3 (CSF-3), GM-CSF (granulocyte-macrophage colony-stimulating factor), IFN alpha (interferon alpha), IFN gamma, IL-1 alpha, IL-1 beta, IL-2, IL-3, IL-4, IL-5, IL-6, IL-7, IL-8 (also known as CXCL8), IL-9, IL-10, IL-12 (also termed as IL-12p70), IL-13, IL-15, IL-16, IL-17A (also termed as CTLA-8), IL-18, IL-20, IL-21, IL-22, IL-23, IL-27, IL-31, LIF (leukemia inhibitory factor), M-CSF (macrophage colony-stimulating factor), MIF (macrophage migration inhibitory factor), TNF alpha, TNF beta, and TSLP (thymic stromal lymphopoietin). The compatible chemokines include BLC (B lymphocyte chemoattractant, also termed as chemokine ligand 13 CXCL13), ENA-78 (epithelial neutrophil-activating protein 78, also known as CXCL5), Eotaxin (also known as CCL11), Eotaxin-2 (also termed as CCL24), Eotaxin-3 (also known as CCL26), Fractalkine (also termed as CX3CL1), Gro-alpha (also known as CXCL1), IP-10 (IFN-γ inducible protein, also termed as CXCL10), I-TAC (IFN-inducible T cell alpha chemoattractant, also known as CXCL11), MCP-1 (monocyte chemoattractant protein-1, also termed as CCL2), MCP-2 ( monocyte chemoattractant protein-2 also known as CCL8), MCP-3 (monocyte chemoattractant protein-3 also known as CCL7), MDC (macrophage-derived chemokine, also termed as CCL22), MIG (monokine induced by interferon-γ, also known as CXCL9), MIP-1 alpha (macrophage inflammatory protein-1 alpha, also termed as CCL3), MIP-1 beta (also known as CCL4), MIP-3 alpha (also termed as CCL20), SDF-1 alpha (stromal cell-derived factor-1 alpha, also known as CXCL12). The compatible growth factors include FGF-2 (fibroblast growth factor-2), HGF (hepatocyte growth factor, MMP-1 (matrix metalloproteinase-1), NGF beta (nerve growth factor-beta), SCF (stem cell factor), and VEGF-A (vascular endothelial growth factor). The compatible soluble receptors are APRIL (a proliferation-inducing ligand), BAFF (B cell-activating factor), CD30, CD40L (also known as CD154), IL-2R (IL-2 receptor, also known as CD25), TNF-RII (tumor necrosis factor receptor II), TRAIL (TNF-related apoptosis-inducing ligand, CD253), and TWEAK (tumor necrosis factor-like weak inducer of apoptosis).

### Determination of serum SDF-alpha and TNF-RII levels in **LRRK2** G2019S KI mice

*LRRK2* G2019S KI mice (RRID: IMSR_JAX:030961) and control wild type (WT) mice (RRID: IMSR_JAX:000664) in the C57BL/6J background were originally purchased from Jackson Laboratory (Bar Harbor, ME). Two batches of mice, aged 3 and 13 months, were used to validate SDF-1 alpha and TNF-RII levels in serum. Both male and female mice were used. Mice were housed in a temperature-controlled room with a 12-h light/dark cycle and had free access to food and water. All procedures were approved by the Institutional Animal Ethical Committee of Massachusetts General Hospital (animal protocol #2018N000039).

Blood was obtained through cardiac puncture and collected in serum separator tubes (BD Vacutainer). Serum samples were analyzed for SDF-1 alpha and TNF-RII levels using mouse SDF-1 alpha and TNF-RII Quantikine ELISA kits (R&D systems, Catalog #: MCX120 and MRT20). ELISA was performed per the manufacturer’s instructions. Recombinant SDF1 and TNF-RII were used as positive controls. f

### Data Processing and Statistical Analysis

The Bioplex Multiplex Immunoassay was performed blinded. The data was collected and sent back to MJFF for unblinding. For data processing, the lower-than-lower limit of detection values was set to 0, while the higher-than-higher limit of detection value was set to 100000. The log2-transformed data was used for all analyses to address outliers and non-normal distributions. Batch effects were investigated using clustered heatmaps and corrected through batch mean centering. For demographic variables, continuous variables (e.g., age) were summarized using means and standard deviations; categorical variables (e.g., sex) were summarized using counts and frequencies. Kruskal-Wallis test and Fisher’s exact test were used to compare the continuous and categorical data, respectively.

A multivariable robust linear regression model, which is less sensitive to outliers, was used to determine associations of analyte levels with *LRRK2* mutations and PD status, conduct comparisons across the different groups (e.g., *LRRK2*+/PD, *LRRK2*+/UC, *LRRK2*-/PD, *LRRK2*-/UC), and assess for any differences while adjusting for potential confounders age, sex, and sample cohort. Correlations were evaluated using the Spearman correlation coefficient. Although the study was exploratory, we assessed p values with and without multiple comparisons adjustment, using Benjamini and Hochberg’s approach, which controls the false discovery rate (FDR). The statistical analyses were conducted using R software (R Development Core Team). All tests were two-sided, and p≤ 0.05 was considered statistically significant. For the preclinical validation of SDF-1 alpha and TNF-RII, one-way ANOVA was employed to analyze differences in *LRRK2* G2019S KI mice and WT controls.

## Results

### GPT-extracted literature overview of cytokines and PD

We performed a GPT-assisted literature review of the 65 cytokines and PD, which includes PD with *LRRK2* mutations. The number of publications discussing cytokines, LRRK2, and PD has increased dramatically in recent years (Supplemental Figure 1). The results for many cytokines appear variable. However, more studies indicated that PD is associated with increased IFN gamma, IL-18, and TNF alpha, among other cytokines, than studies reporting no associations. This was observed in the “all papers” search (Figure 1), including all studies across various hosts, including humans, animals, and cell lines (Figure 1A), and in human studies only (Figure 1B). Our “primary paper” search, excluding reviews, editorials, letters, and pre-prints, showed similar results (Supplemental Figure 2). Our GPT-assisted search did not separate blood and CSF.

**Figure 1.**
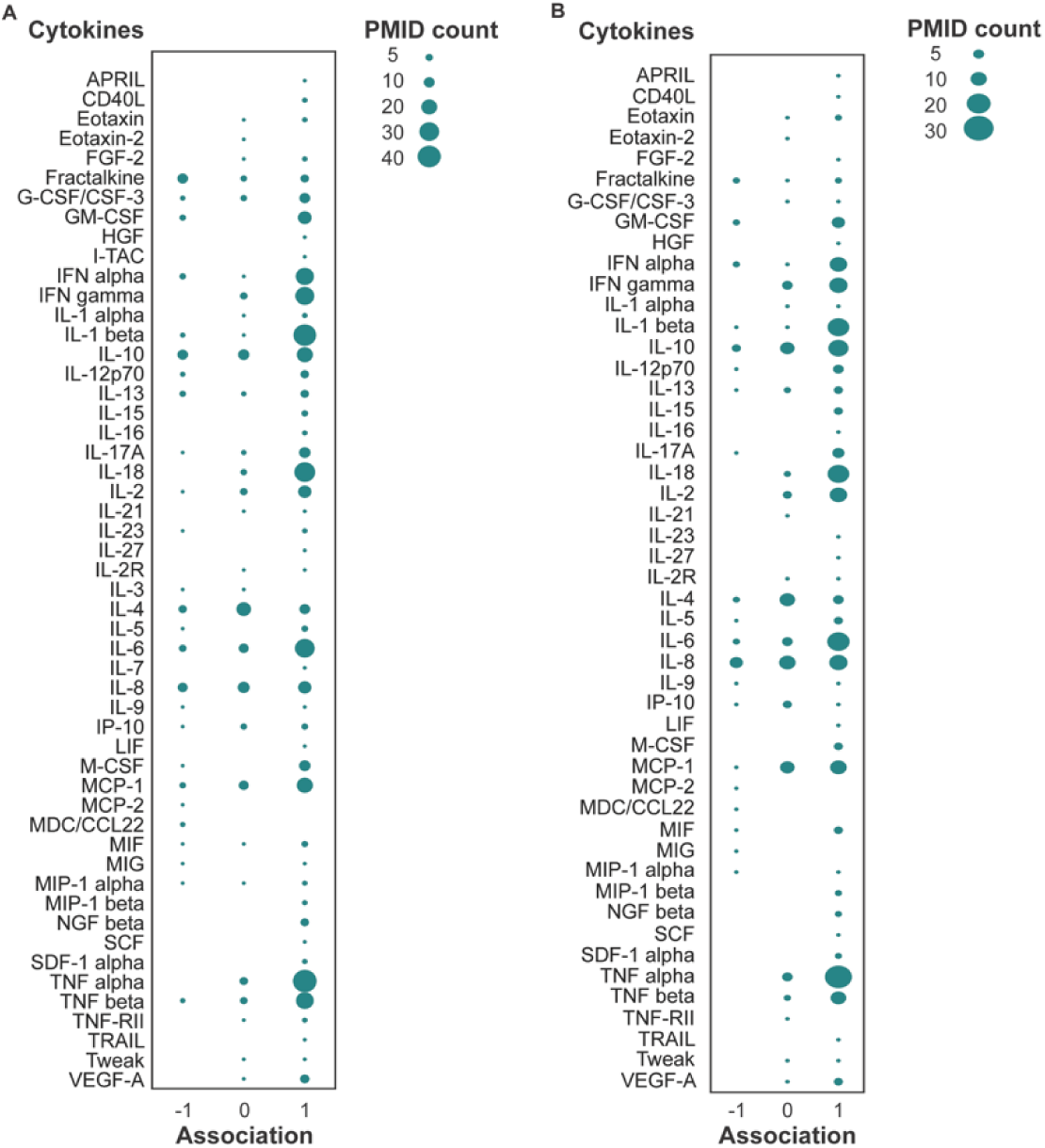
GPT-extracted literature overview of cytokines and PD. Scores of 1 were given to positive associations between higher cytokine concentrations and PD, −1 to negative associations between lower cytokine concentrations and PD, and 0 to no associations. Bubble size represents the number of papers supporting the association scores. A: Studies across various hosts, such as humans, animals, and cell lines B: Human studies only. A total of 1060 papers were extracted and filtered for associations with PD or LRRK2, and then a total of 401 papers were used for analysis.

### Characteristics of the Participants

Serum analytes were collected from a total of 651 subjects available in the LCC (Table 1A) with an approximate 2:1 ratio in the number of *LRRK2*+ subjects to the number of *LRRK2*-subjects and an approximate 1:2 ratio in the number of PD subjects to the number of non-PD subjects. PD subjects were significantly older than those without a PD diagnosis (p<0.001). There was also a statistically significant difference (p=0.022) in the male-to-female ratios in each group, as there was a greater proportion of females than males in every group except the *LRRK2-/*PD group. Among the 129 subjects who provided CSF, there was a nearly 1:1 ratio in the number of PD subjects to the number of UC and in the number of *LRRK2*+ subjects to those without a *LRRK2* mutation. There was a statistically significant age difference (p=0.024) among the groups, with the *LRRK2+/*PD group having the oldest subjects compared to the other three groups. There was no significant difference in terms of sex throughout the groups (Table 1B). Table 1C comprises those who had both serum and CSF samples. There was a near 1:1 ratio between PD: UC and *LRRK2+*:*LRRK2-* subjects, and there was no significant difference in the mean age and sex across the four groups in this subset (Table 1C).

**Table 1.**
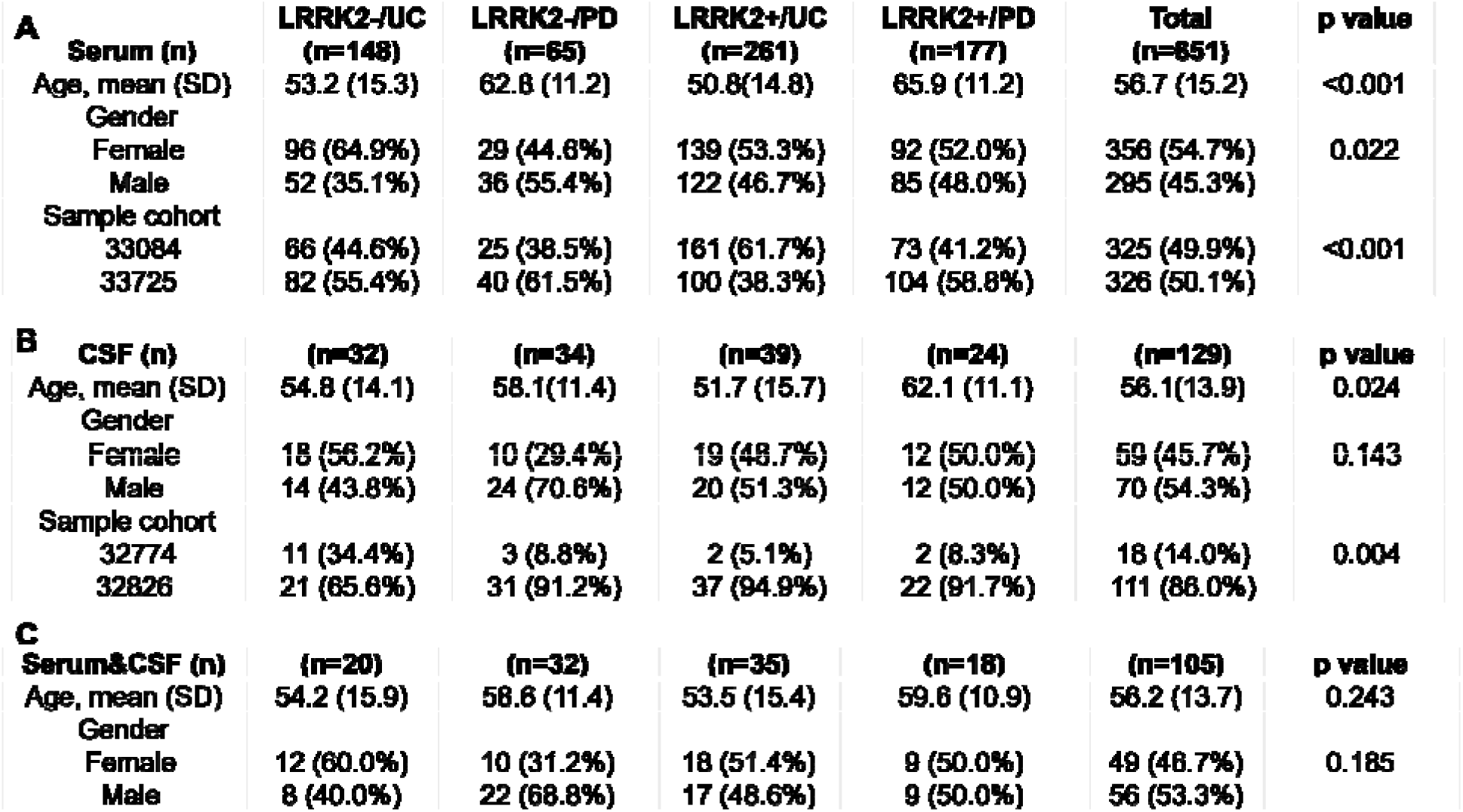
Demographics and features by *LRRK2* and PD status of participants contributing analyzed samples. Sample size, mean age (SD), sex, and p value by group status for (A) serum, (B) CSF, and (C) matching serum and CS samples. A and B also list group status according to sample cohort.

### Presence of soluble immune factors in serum and CSF samples

A total of 65 analytes were quantified from the four groups of serum and CSF samples. In serum, 22 out of the 65 analytes were detectable in over 50% of the samples. These analytes included APRIL, BLC, CD30, ENA-78, Eotaxin, Eotaxin-2, HGF, IL-16, IL-18, IL-2R, IL-7, IP-10, MCP-1, MCP-2, MDC/CCL22, MIF, MMP-1, SCF, SDF-1 alpha, TNF-RII, Tweak, and VEGF-A. In contrast, 63 analytes were detectable in more than 50% of the CSF samples, except for IL-5 and TRAIL, which have detection rates of 49.6% and 36.4%, respectively, in CSF samples. The means of tested analytes and their detection rates in percentages in serum and CSF samples were summarized in Figure 2. A broad range of soluble immune factors in CSF supports the potential involvement of CNS immune responses in PD. It provides a rationale for identifying differentially expressed analytes in subjects with PD and *LRRK2* mutations.

**Figure 2.**
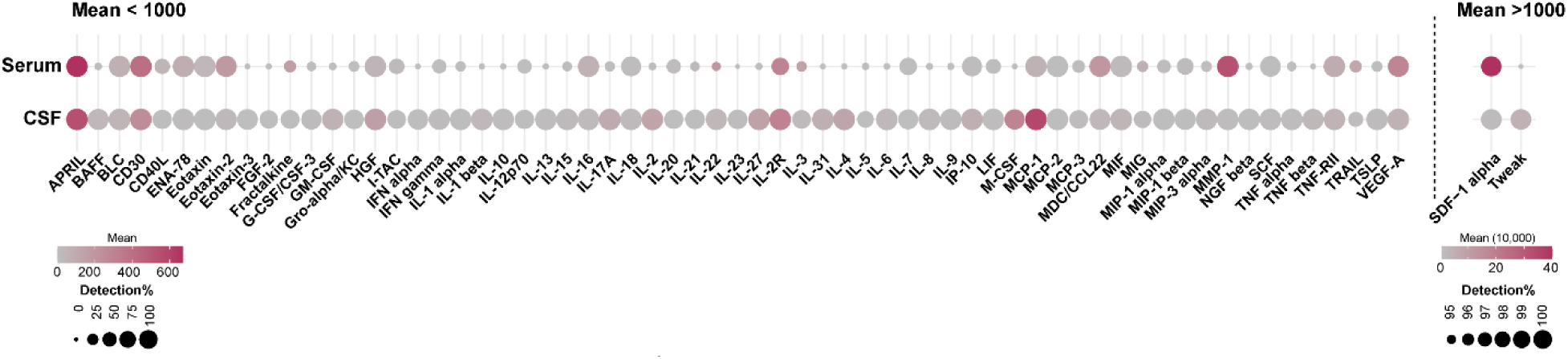
Means of log_2_ raw concentration (pg/mL) of soluble immune factors and their detection percentage in serum and CSF samples of the LCC.

### Differentiated soluble immune factors in serum and CSF in LRRK2 mutation carriers

To identify changes in soluble immune regulators associated with *LRRK2* mutations, we compared serum and CSF results from subjects carrying *LRRK2* mutations (n=438) with those from non-carriers (n=213), irrespective of PD status. After adjusting for age, sex, sample cohort, and PD status, multivariable linear regression analysis identified seven elevated and two reduced immune factors with p-value <0.05 in the *LRRK2* carrier group based on (Figure 3A). Notably, *LRRK2* mutation carriers demonstrated an increase in SDF-1 alpha, a stromal cell-derived chemokine in serum, compared to non-carriers (p=0.0007). The difference is statistically significant after adjusting for multiple comparisons (p adj=0.026) (Figure 3B). Another major immune regulator, TNF-RII, was found to have a higher concentration in *LRRK2* mutation carriers compared to non-carriers, and a significant difference was observed between the two groups before and after multiple comparison adjustments (p=0.0008, p adj=0.026) (Figure 3C). Compared to non-carriers, additional elevated analytes in serum of *LRRK2* mutation carriers included VEGF-A (p=0.002), MIP-1 beta (p=0.003), MCP-1 (p=0.015), MIF (p=0.015), and IP-10 (p=0.024). In contrast, IL-20 (p=0.013), LIF (p=0.038), and IL-7 (p=0.048) were reduced in the serum of *LRRK2* mutation carriers compared to non-carriers (Figure 3D). However, p values for these analytes were over 0.05 after multiple comparisons adjustment.

**Figure 3.**
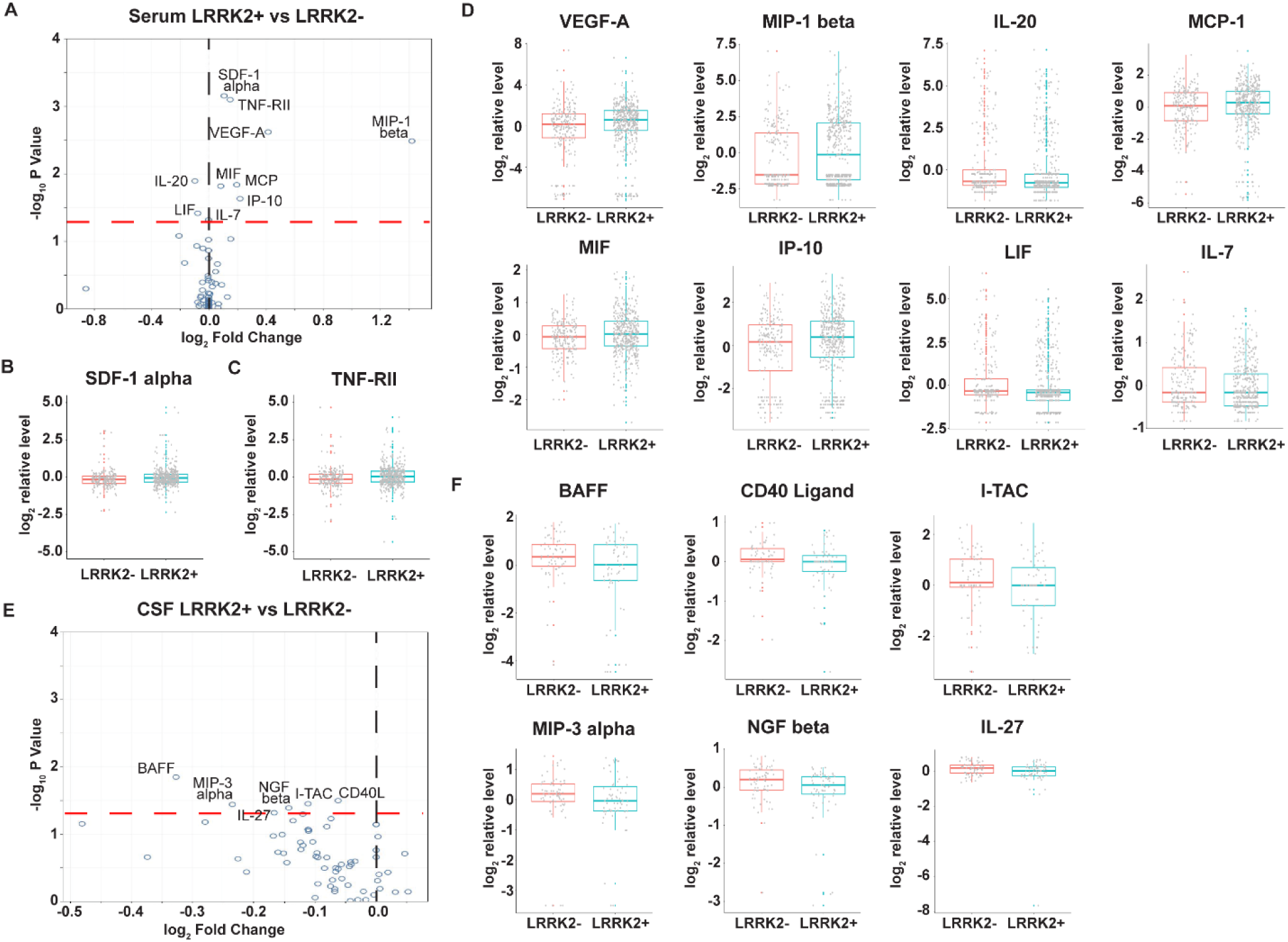
Changes of soluble immune factors in serum and CSF samples of LCC participants carrying *LRRK* mutations. A: Volcano plot of serum analytes comparing *LRRK2*+ (n=438) and *LRRK2*-(n=213) participants. B-D: Batch-corrected concentrations of SDF-1 alpha, TNF-RII, VEGF-A, MIP-1 beta, MCP-1, MIF, IP-10, IL-20, LIF, and IL-in serum of *LRRK2*+ and *LRRK2*-participants. E: Volcano plot of CSF analytes comparing *LRRK2*+ (n=63) vs *LRRK2*-(n=66) precipitants. F: Batch-corrected concentrations of BAFF, CD40-Ligand, I-TAC, MIP-3 alpha, NGF beta, and IL-27 in CSF comparing *LRRK2*+ and *LRRK2*-participants. The volcano plots illustrate the data normalized to the mean. The black and red dashed line represents “log_2_ Fold Change” of 0 and “-log_10_ P Value” of 1.3, respectively. Th boxplots illustrate the median (the horizontal line within the box), and the box represents the upper and lower quartiles. Robust linear regression adjusting for age, sex, sample cohort, and PD status.

In contrast to serum, *LRRK2* mutations were only associated with moderately reduced soluble immune markers in **CSF**. Analysis of 129 samples revealed six reduced analytes in subjects carrying *LRRK2* mutations (n=63) compared with those non-carriers (n=66), irrespective of PD status (Figure 3E). These analytes, which included BAFF (p=0.014), CD40-Ligand (p=0.032), I-TAC (p=0.035), MIP-3 alpha (p=0.036), NGF beta (p=0.041), and IL-27 (p=0.048), were all reduced (Figure 3F). However, none of these differences remained statistically significant after adjusting for multiple comparisons.

We calculated **CSF: serum ratios** of all analytes from the **subset** of matching 105 serum and CSF samples, which had well-balanced numbers of *LRRK2* mutation carriers (n=53) vs non-carrier (n=52). CSF: serum ratios for TNF-RII and SDF-1 alpha were lower in *LRRK2* carriers than non-carriers (p=0.005 and 0.007, respectively), which were most likely driven by changes in the levels of the two analytes in serum. Additionally, CSF: serum ratio for APRIL, another member of the TNF superfamily, was lower in *LRRK2* carriers than non-carriers in this subset (p=0.031) (Figure 4).

**Figure 4.**
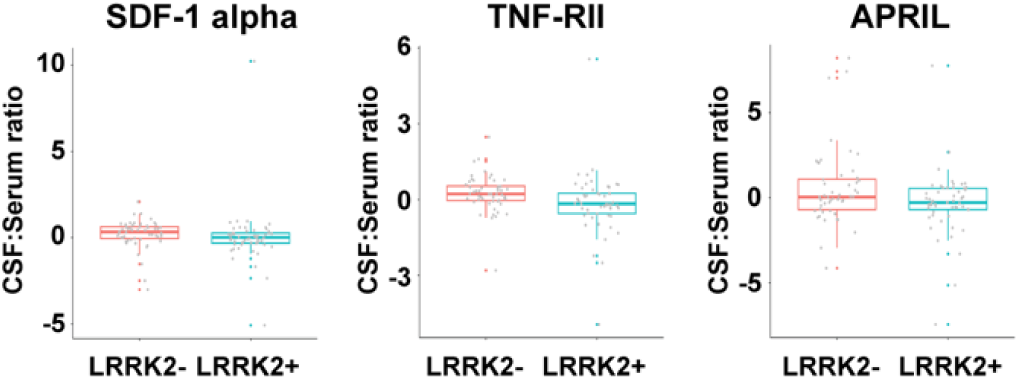
Altered CSF: serum ratios of soluble immune factors comparing *LRRK2*+ and *LRRK2*-participants. CSF: serum ratios for SDF-1 alpha, TNF-RII, and APRIL in *LRRK2*+ (n=53) and *LRRK2*-(n=52) participants. The boxplots illustrate the median (using a horizontal line within the box) and the box represents the upper and lower quartiles. Robust linear regression adjusting for age, sex, sample cohort, and PD status.

### Differentiated soluble immune factors in serum and CSF in PD subjects

We compared serum and CSF results from PD (n=182) and control subjects (n=409), irrespective of *LRRK2* mutations. Surprisingly, there were no significant differences in serum concentration of all the analytes except marginally lower SCF in PD than in the control group (p=0.045) (Figure 5A&B). In CSF, no analytes were elevated. Concentrations of MIF (p=0.002), MMP-1 (p=0.005), CD30 (p=0.030), Tweak (p=0.040), and SDF-1 alpha (p=0.042) were lower in PD (n=58) as compared to control subjects (n=71) (Figure 5C&D). However, none of these difference remained statistically significant after adjusting for multiple comparisons.

**Figure 5.**
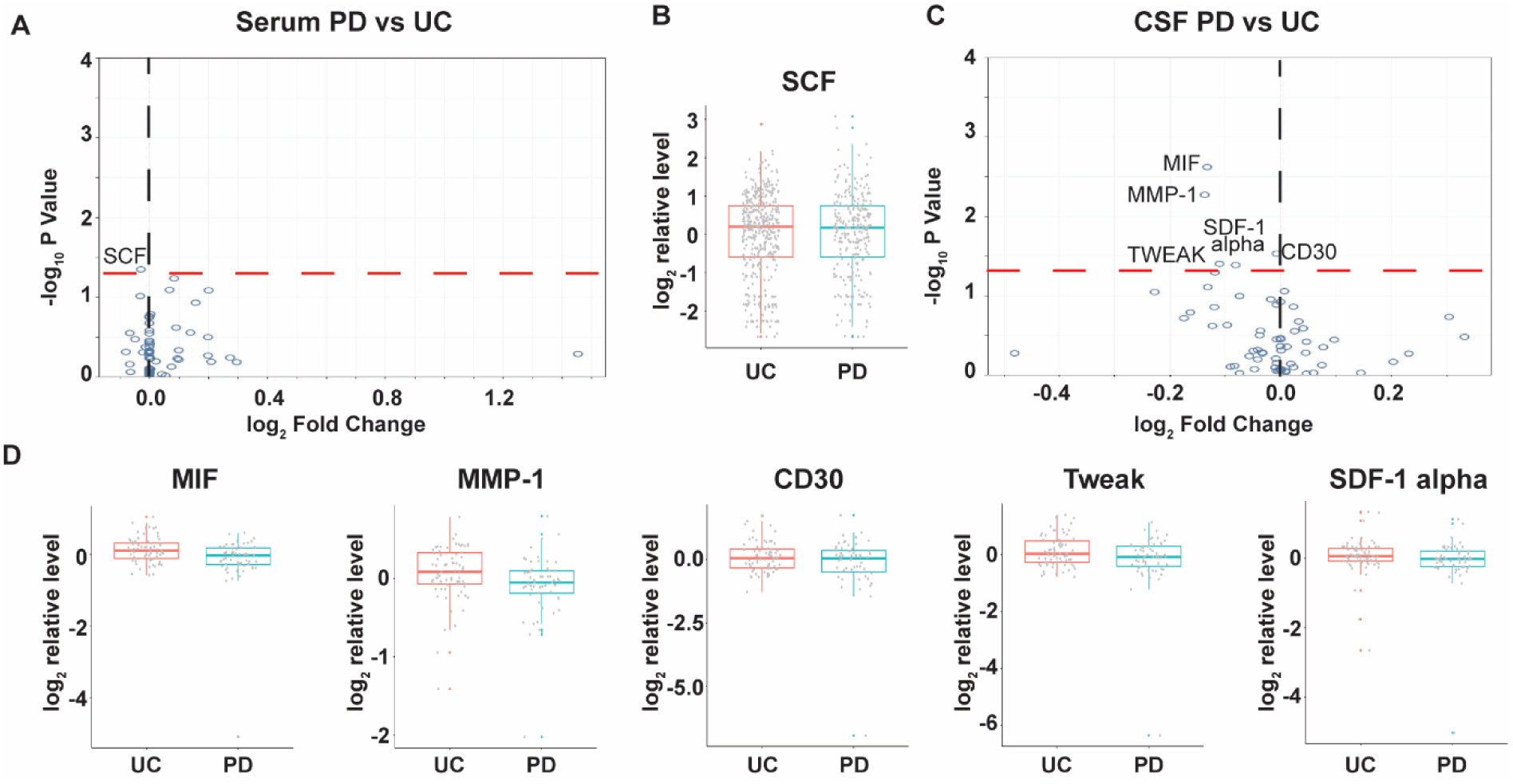
Changes of soluble immune factors in serum and CSF samples of LCC participants with PD. A: Volcano plot of serum analytes comparing PD (n=182) and unaffected controls (n=409). B: Batch-corrected concentrations of SCF in serum of PD subjects and unaffected controls. C: Volcano plot of CSF analytes PD (n=58) vs unaffected controls (n=71). D: Batch-corrected concentrations of MIF, MMP-1, CD30, Tweak, and SDF-1 alpha in CSF of PD subjects and unaffected controls. The volcano plots illustrate the data normalized to the mean. The black and red dashed line represents “log_2_ Fold Change” of 0 and “-log_10_ P Value” of 1.3, respectively. The boxplots illustrate th median (the horizontal line within the box), and the box represents the upper and lower quartiles. Robust linear regression adjusting for age, sex, sample cohort, and *LRRK2* mutation.

In PD subjects from the subset with matching serum and CSF, the CSF: serum ratios for CD30 (p=0.002), MCP-2 (p=0.010), and APRIL (p=0.021) were lower, while Eotaxin was higher (p=0.030) compared to the control (Figure 6).

**Figure 6.**
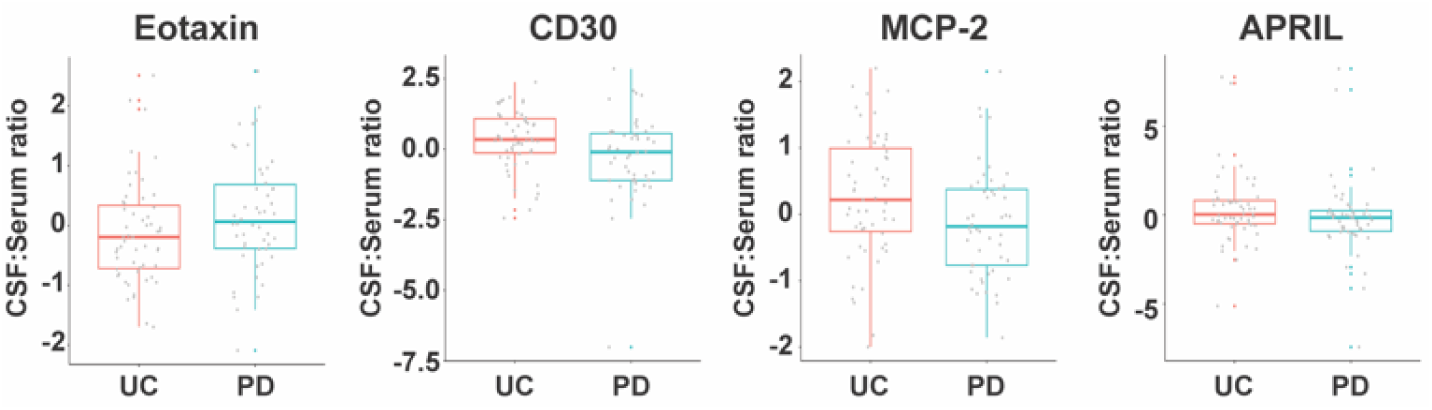
Altered CSF: serum ratios of soluble immune factors in PD subjects compared with unaffected controls. CSF: serum ratios for Eotaxin, CD30, MCP-2, and APRIL in PD subjects (n=50) and UC subjects (n=55). The boxplots illustrate the median (using a horizontal line within th box) and the box represents the upper and lower quartiles. Robust linear regression adjusting for age, sex, sampl cohort, and *LRRK2* mutation.

### *LRRK2* PD-associated changes in immune markers

To identify possible inflammatory and immune markers related to *LRRK2* and PD, we performed four-group comparisons using the robust linear regression models adjusting for age, sex, and sample cohort. In serum, compared with idiopathic PD (*LRRK2-*/PD), the *LRRK2+*/PD group had lower CD30 (p=0.020). Compared with *LRRK2* carriers without PD (*LRRK2+*/UC), *LRRK2+*/PD groups had lower SCF (p=0.025) (Table 2A). There were no significant differences in serum concentrations of SDF-1 alpha and TNF-RII between PD subjects with LRRK2 mutations and those without, even though a trend for increased concentrations was observed for both. The overall difference in SDF-1 alpha between *LRRK2*+ and *LRRK2*-groups (effect size=0.1135) in Fig 3 appears to be primarily attributable to the difference between *LRRK2*+ and *LRRK2*-subjects within the UC group (effect size=0.0416, p=0.0008). These results support that the difference in SDF-1 alpha is predominantly driven by the UC subgroup. Additionally, we did not observe any significant differences in serum concentrations of SDF-1 alpha and TNF-RII between *LRRK2* carriers with PD and those without (Table 2A).

**Table 2.**
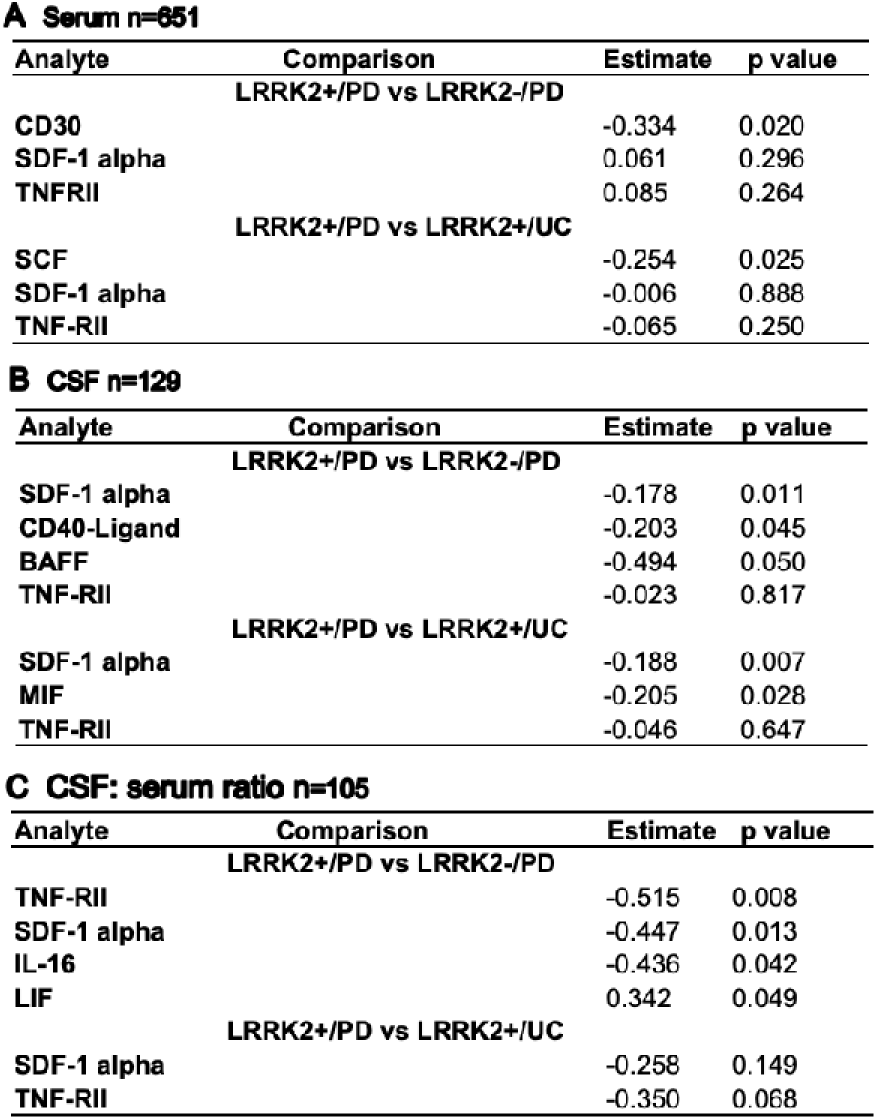
Altered analytes in *LRRK2*+PD subjects from 4-group comparisons. A: Full set of serum samples. B: Full set of CSF samples. C: CSF:serum ratios. Robust linear regression adjusting for age, sex, and sample cohort.

A Serum n=651
| Analyte | Comparison | Estimate | p value |
| --- | --- | --- | --- |
| **LRRK2+/PD vs LRRK2-/PD** |  |  |  |
| CD30 |  | -0.334 | 0.020 |
| SDF-1 alpha |  | 0.061 | 0.296 |
| TNFRII |  | 0.085 | 0.264 |
| **LRRK2+/PD vs LRRK2+/UC** |  |  |  |
| SCF |  | -0.254 | 0.025 |
| SDF-1 alpha |  | -0.006 | 0.888 |
| TNF-RII |  | -0.065 | 0.250 |

B CSF n=129
| Analyte | Comparison | Estimate | p value |
| --- | --- | --- | --- |
| **LRRK2+/PD vs LRRK2-/PD** |  |  |  |
| SDF-1 alpha |  | -0.178 | 0.011 |
| CD40-Ligand |  | -0.203 | 0.045 |
| BAFF |  | -0.494 | 0.050 |
| TNF-RII |  | -0.023 | 0.817 |
| **LRRK2+/PD vs LRRK2+/UC** |  |  |  |
| SDF-1 alpha |  | -0.188 | 0.007 |
| MIF |  | -0.205 | 0.028 |
| TNF-RII |  | -0.046 | 0.647 |

| Analyte | Comparison | Estimate | p value |
| --- | --- | --- | --- |
| **LRRK2+/PD vs LRRK2-/PD** |  |  |  |
| TNF-RII |  | -0.515 | 0.008 |
| SDF-1 alpha |  | -0.447 | 0.013 |
| IL-16 |  | -0.436 | 0.042 |
| LIF |  | 0.342 | 0.049 |
| **LRRK2+/PD vs LRRK2+/UC** |  |  |  |
| SDF-1 alpha |  | -0.258 | 0.149 |
| TNF-RII |  | -0.350 | 0.068 |

In CSF, compared to *LRRK2*-/PD, the *LRRK2*+/PD group had lower SDF-1 alpha (p=0.011), which is the reverse of what was observed in the serum. CD40-Ligand (p=0.045) and BAFF (p=0.050) were also lower in the *LRRK2*+/PD group (Table 2B). SDF-1 alpha (p=0.007) was lower, in addition to MIF (p=0.028), in the *LRRK2+*/PD group compared to the *LRRK2*+/UC group (Table 2B). The difference of TNF-RII was in the opposite direction in CSF to that in serum, comparing the *LRRK2+*/PD group to the *LRRK2*-/PD group, but the magnitude of the difference was minor (Table 2B). These results are overall consistent with the trends observed in the *LRRK2*+ vs *LRRK2*- and PD vs UC groups.

In terms of the CSF: serum ratios of the analytes, the *LRRK2+*/PD group had significantly decreased concentration ratios of TNF-RII (p=0.008), SDF-1 alpha (p=0.013), and IL-16 (p=0.042) compared to *LRRK2*-/PD. No significant changes were observed between the *LRRK2*+/PD and *LRRK2*+/UC groups in concentration ratios of TNF-RII, SDF-1 alpha, or other analytes (Table 2C). None of the analyte differences in the four-group comparisons remained statistically significant after adjusting for multiple comparisons.

### Correlations between serum and CSF soluble immune factors

Given that the altered analytes in serum and CSF related to *LRRK2* mutations and PD were largely non-overlapping or had an inverse relationship, we analyzed the correlation between serum analytes and CSF analytes in the subset of matching serum and CSF samples using the Spearman correlation coefficient. Four out of 65 analytes showed a statistically significant correlation between serum and CSF. As shown in Figure 7, SCF, IP-10, and Eotaxin-2 in serum and CSF were positively correlated (p=0.0007, 0.001 and 0.033, respectively). Correlations of SCF and IP-10 between serum and CSF were still significant after multiple comparison adjustments (p adj=0.031 for both). Serum and CSF BAFF concentrations were negatively correlated (p=0.01).

**Figure 7.**
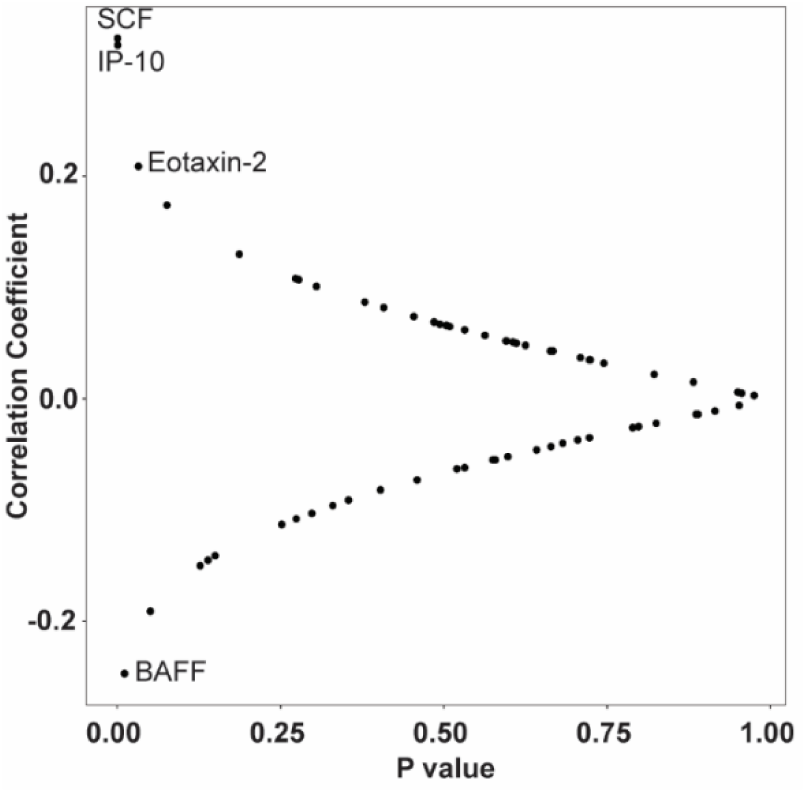
Correlation between serum and CSF concentrations o f soluble immune factors. n=105. Spearman correlation coefficient test.

### Serum levels of SDF-1 alpha and TNF-RII in *LRRK2* G2019S KI mice

To validate our findings of serum SDF-1 alpha and TNF-RII in *LRRK2* mutation carriers, we assessed by ELISA the levels of SDF-1 alpha and TNF-RII levels in serum samples from *LRRK2* G2019S KI mice and age-matched WT control mice. At 3 months of age, there was no significant difference in either serum SDF-1 alpha or TNF-RII between *LRRK2* G2019S KI mice and WT controls (Figure 8A&B). However, serum SDF-1 alpha levels were significantly higher in the *LRRK2* G2019S KI group compared to WT at the age of 13 months old (*p*=0.016). Higher serum TNF-RII concentrations were also observed in older *LRRK2* G2019S KI mice compared to WT controls, though not statistically significant (Figure 8A&B). These results indicate that the *LRRK2* mutation is associated with an increase in serum SDF-1 alpha, similar to what is observed in human *LRRK2* carriers, who had a mean age of 57. Non-carriers had a mean age of 56.

**Figure 8.**
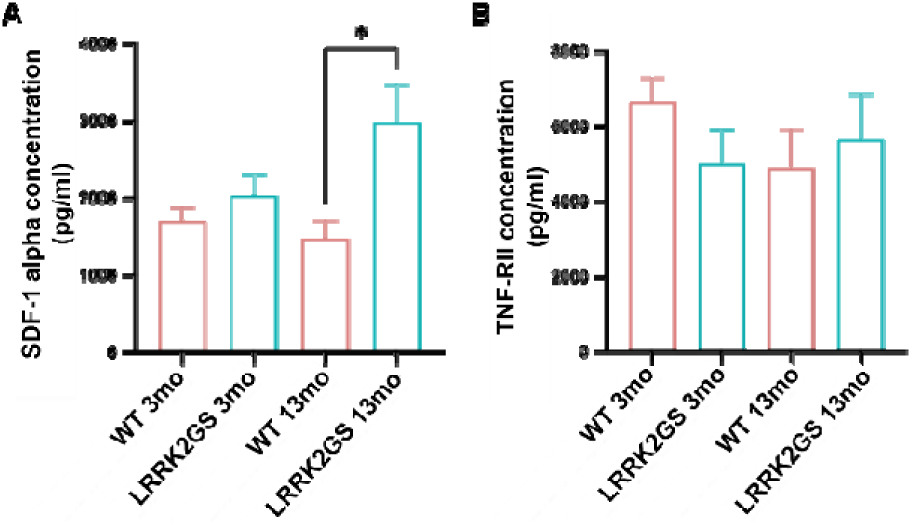
Serum levels of SDF-1 alpha and TNF-RII i LRKK2 G2019S KI mice. . Age- and sex-matched LRKK G2019S KI (*LRRK2*GS) and control WT mice at the ages of months (3mo) and 13 months (13mo) were used, and Serum levels of SDF-1 alpha and TNF-RII were determine by ELISA. A: Serum levels of SDF-1 alpha. B: Serum levels of TNF-RII. Data are presented as mean ± SEM. One-way ANOVA followed by Tukey’s test. *p@0.05. n=6 (3 males and 3 females) all groups.

## Discussion

Our study using the LCC as a discovery cohort revealed that *LRRK2* mutations were associated with increased immune markers in serum, not CSF, whereas PD was associated with decreased immune markers in CSF, not serum.

Specifically, subjects harboring pathogenic mutations in the *LRRK2* gene show increased production of immune regulators in serum. SDF-1 alpha and TNF-RII, in particular, have been identified as potential serum biomarkers for *LRRK2* mutation carriers, though not to the extent of differentiating *LRRK2+* /PD from *LRRK2-*/PD or *LRRK2+*/UC from *LRRK2-*/UC. The increased serum levels of SDF-1 alpha related to *LRRK2* mutation were validated in *LRRK2* G2019S KI mice at an older age.

SDF-1 alpha, also known as CXCL12, is a potent chemotactic protein produced by bone marrow stromal cell lines. Through its two receptors, CXCR4 and CXCR7, SDF-1 alpha acts as a key homeostatic chemokine regulating embryogenesis, hematopoiesis, and angiogenesis.^24^ It promotes migration, proliferation, and maturation of hematopoietic progenitor cells, endothelial cells, and leukocytes. SDF-1 alpha expression is elicited in primary and secondary lymphoid organs as a part of the homeostatic mechanism regulating immune cell development and trafficking.^24^ It is also involved in the regulation of inflammation with a vital role in wound healing and tissue repair in inflammatory diseases.^25^ SDF-1 alpha has been reported to promote cancer arthritic diseases but has also been shown to be cardioprotective by promoting stem cell homing, angiogenesis, and remote ischemic conditioning.^26^ In the CNS, the SDF-1 alpha signaling promotes the proliferation, differentiation, and migration of neural precursor cells and mediates axonal elongation and branching after cerebral ischemia.^27^ SDF-1 alpha can promote microglial phagocytosis of amyloid beta and be neuroprotective in Alzheimer’s disease (AD).^28^ More recent studies have shown that SDF-1 alpha may also be proinflammatory and mediates alpha-synuclein-induced microglia accumulation.^29^ We could not find evidence of serum SDF-1 alpha in *LRRK2* mutation carriers in the literature. A study showed a higher level of SDF-1 alpha in CSF of asymptomatic *LRRK2* mutation carriers but statistically not significant.^29^ In PD patients, blood SDF-1 alpha was found to be significantly higher than control subjects.^30,31^ An upregulation of the CXCL12/CXCL4 signaling pathway has been shown to be involved in the loss of dopaminergic neurons in animal models.^32^ Whether increased SDF-1 alpha is mechanistically involved in *LRRK2* mutations and increased PD risk requires more investigations.

TNF alpha has been reported to be significantly higher in CSF and serum of asymptomatic *LRRK2* mutation carriers compared to healthy controls.^16,33^ TNF alpha binds to TNF RI and TNF-RII and plays a crucial role in innate and adaptive immune responses. While TNF-RII is ubiquitously expressed in almost all cell types and predominantly mediates proinflammatory responses, TNF-RII has been shown to express predominantly on regulatory T cells (Tregs).^34,35^ Through the immune modulatory function of Tregs, TNF-RII promotes tissue homeostasis and regeneration.^36,37^ In a mouse model of AD, activation of TNF-RII was found to mitigate cognitive deficits and neuropathology through decreasing amyloid β production and increasing its clearance by glial cells.^38^ Other studies have also reported the activation of TNF-RII to be neuroprotective in AD.^39,40^ Conversely, significant deficiency in TNF-RII signaling has been demonstrated in various autoimmune diseases.^41,42^ Soluble TNF receptors are released into biological fluids by proteolytic cleavage of the transmembrane receptors. Soluble TNF-RII binds to and effectively neutralizes TNF alpha. It acts as a negative regulator of TNF-alpha signaling, exerting anti-inflammatory effects and promoting tissue homeostasis and regeneration.^35,36^ Levels of soluble TNF-RII are elevated in the serum of patients with multiple sclerosis.^43^ TNF-RII has been increasingly recognized as a potential therapeutic target for various inflammatory conditions, including rheumatoid arthritis and inflammatory bowel disease.^44,45^ Possible impact of elevated serum TNF-RII in *LRRK2* carriers should be further investigated.

Other *LRRK2* mutations-related changes in serum include increases in VEGF-A, which promotes angiogenesis and growth of solid tumors, MIP-1 beta, which is produced mainly by macrophages and monocytes and has been shown to orchestrate protective responses against various viral infections,^46,47^ MCP-1, which promotes the migration of inflammatory cells and is a key protein in tumor development,^48^ MIF, which sustains the survival and function of macrophages and drives inflammation,^49,50^ and IP-10, which controls cell growth and development including tumor cells growth and angiostasis.^51^ Among the reduced serum analytes, IL-20 induces the proliferation of epithelial cells and the production of proinflammatory factors.^52^ LIF is a pleiotropic cytokine that maintains the homeostasis and regeneration of multiple tissues.^53^ IL-7 is crucial for B and T cell development.^54^

Compared with changes identified in serum, *LRRK2* mutations are associated with a completely different profile of immune and inflammatory factors in CSF, which showed moderately reduced signals related to the regulation of B cell selection and survival (BAFF),^55^ T-B cell communications (CD40-Ligand through interaction with CD40),^56^ immune cell recruitment (I-TAC), dendritic cell trafficking (MIP-3 alpha), neural development and survival (NGF beta), and inhibition of inflammation (IL-27).^57^ Many of these cytokines and chemokines have multifaceted functions and can be tissue- and cell-dependent.

The general reduction of inflammatory and immune markers in CSF and the overall null findings in the serum of PD subjects in this large cohort of samples was unexpected. Out of 65 immune markers analyzed, only SCF was marginally reduced in serum samples from PD subjects compared to those from healthy controls. SCF is a survival and growth factor for hematopoietic stem and progenitor cells. Previous studies have shown increased or decreased levels of SCF in PD.^31,58^ The change in SCF in our study was mild and non-significant after multiple comparison adjustments.

The reduced signals in CSF include MIF, MMP-1, which is involved in extracellular matrix remodeling and regulating proinflammatory cytokines,^59^ CD30, which belongs to the TNF receptor family and mediates pro-survival signal,^33^ Tweak, which is also a TNF superfamily member and can stimulate inflammatory cytokines and determine synaptic function,^60^ and SDF-1 alpha. None of these analytes showed statistical significance after multiple comparison adjustments. Literature evidence is limited for these immune markers in CSF of PD patients. Previous studies using serum examples have found significantly lower levels of MMP-1,^61^ and higher levels of MIF in PD patients.^62^ The exact roles of these immune regulators in PD have yet to be elucidated.

The non-overlapping patterns between changed analytes in serum and CSF comparing either *LRRK2+* vs *LRRK2-* or PD vs UC revealed overall disconnection between serum and CSF in the LCC. This is supported by correlation analysis showing no positive correlations between serum and CSF for all but two of the analytes after multiple comparison adjustment (SCF and IP-10). The disconnection suggests that these analytes may be produced in different compartments and that the integrity of the blood-brain barrier (BBB) may not be affected by either *LRRK2* mutations or PD. Active transport mechanisms across the BBB may also be involved in maintaining the differences between serum and CSF. Additionally, differences in the kinetics of cytokine and chemokine production and clearance between serum and CSF may result in discrepancies in their levels at the time of sampling.

Interestingly, while there are no statistically significant differences in APRIL levels in either serum or CSF among the four groups of subjects stratified based on *LRRK2* mutation and PD status, reduced CSF: serum ratio of APRIL was significantly associated with both *LRRK2* mutations and PD. APRIL, also known as tumor necrosis factor ligand superfamily member 13 (TNFSF13), is a key molecule in regulating B cell survival, maturation, and differentiation.^63^ It promotes the production of immunoglobulins and supports long-term humoral immunity. A previous study has shown that APRIL enhances midbrain dopaminergic axon growth and nigrostriatal projection.^64^ Dysregulation of BAFF/APRIL has been implicated in autoimmune disease, cancer, as well as PD.^65^ Together with reduced levels of two other TNF superfamily members, BAFF and CD40L, our findings may suggest suppressed central relatively to peripheral regulating network for B cell proliferation, survival, and antibody production, particularly in *LRRK2* carriers.

Our study evaluated biomarkers associated with *LRRK2* and PD in one of the most extensive cohorts. It provided comprehensive immune and inflammatory profiles associated with *LRRK2* mutations and PD in blood and CSF. The results revealed that *LRRK2* mutations are associated with increases in regulatory cytokines in serum, particularly the potential of SDF-1 alpha and TNF-RII as serum biomarkers for *LRRK2* mutation carriers. Those with clinically diagnosed PD, regardless of etiology, did not show strong signals in serum but reduced proinflammatory analytes in CSF. These results suggest that while PD mostly shows impaired central immune responses, *LRRK2* mutations may be associated with enhanced peripheral immune regulatory functions. Numerous studies have demonstrated elevated levels of various immune markers linked to either LRRK2 or PD, whereas some reports indicate inconsistencies in immune marker expression.^8,9,10,11,12,16,20,66^ This highlights the complexity of immune and inflammatory-related responses associated with LRRK2 and PD. Our study did not find changes in TNF alpha, IL-6, IL-1 beta, IL-10, IFN gamma, or IL-18 in either serum or CSF, comparing *LRRK2* or PD status. The discrepancies across studies could stem from various factors, including clinical heterogeneity among participants, variations in inclusion criteria, differences in sample sizes, and inconsistencies in sample processing or analytical techniques. Additionally, the timing of sample collection, differences in disease stage or progression among subjects, and potential environmental or other genetic factors may also contribute to the mixed results from different studies.

Although we have adjusted for relevant covariates across groups in our analyses, including age and sex, and PD status or LRRK2 mutation when applicable, unmeasured confounders may influence our results, such as PD medications, disease duration, sampling timing, and sample storage duration. Other limitations inherited from the LCC include possible misclassification of PD and UC, given that there was no pathological diagnosis. Further, sample selection bias cannot be excluded. Future studies should first validate these findings in an independent cohort. Additionally, functional valuation studies should be conducted to explore mechanistic associations between candidate markers and *LRRK2* mutations and/or PD, especially for SDF-1 alpha and TNF-RII.

## Data Availability Statement

Data used in the preparation of this article is openly available to qualified researchers. Demographic data was obtained from the LRRK2 Cohort Consortium Database (www.michaeljfox.org/data-sets). All other resources and codes used, including raw data and information about all the publications related to Figure 1, in this study can be found at DOI:10.5281/zenodo.14861830.

## Acknowledgments

This research was funded by Aligning Science Across Parkinson’s grant ASAP-000312 through the Michael J. Fox Foundation for Parkinson’s Research (MJFF) and NIH grant R01NS102735. The funders had no role in the study design, data collection, analysis, or decision to publish the manuscript. For open access, the author has applied a CC-BY public copyright license to the Author Accepted Manuscript (AAM) version arising from this submission. Data and biospecimens used in this article were obtained from the MJFF-sponsored LRRK2 Cohort Consortium (LCC) under LCC project ID 514. For up-to-date information on the study, visit www.michaeljfox.org/lcc. The LCC is coordinated and funded by MJFF. The authors thank the participants, investigators, and staff of the LCC.

---

**Supplemental Figure 1.**
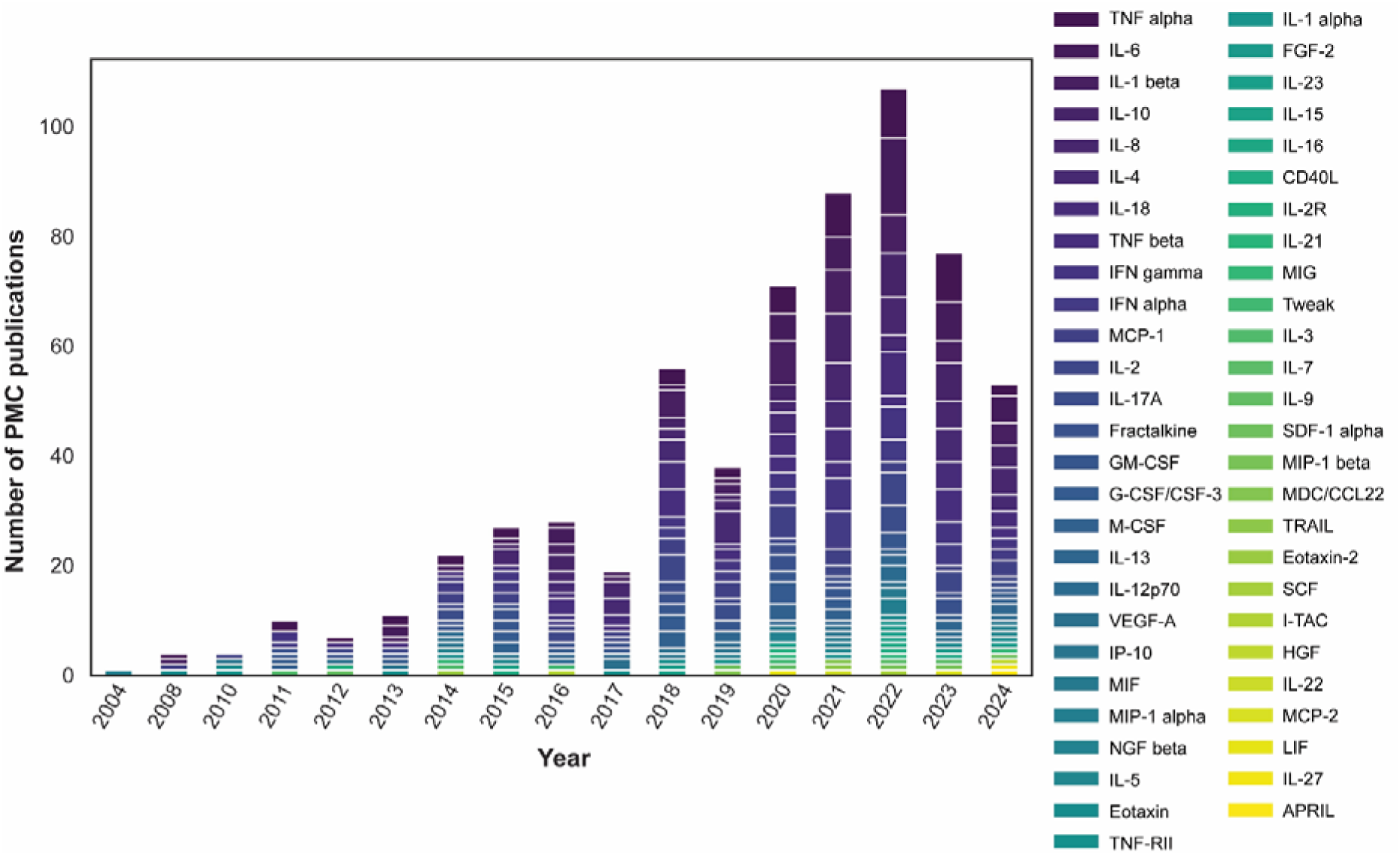
The number of PubMed Central (PMC) publications discussing cytokines, LRRK2, and Parkinson’s disease by publication year. A total of 401 unique papers were retrieved from PubMed and analyzed for 53 cytokines based on keyword search.

**Supplemental Figure 2.**
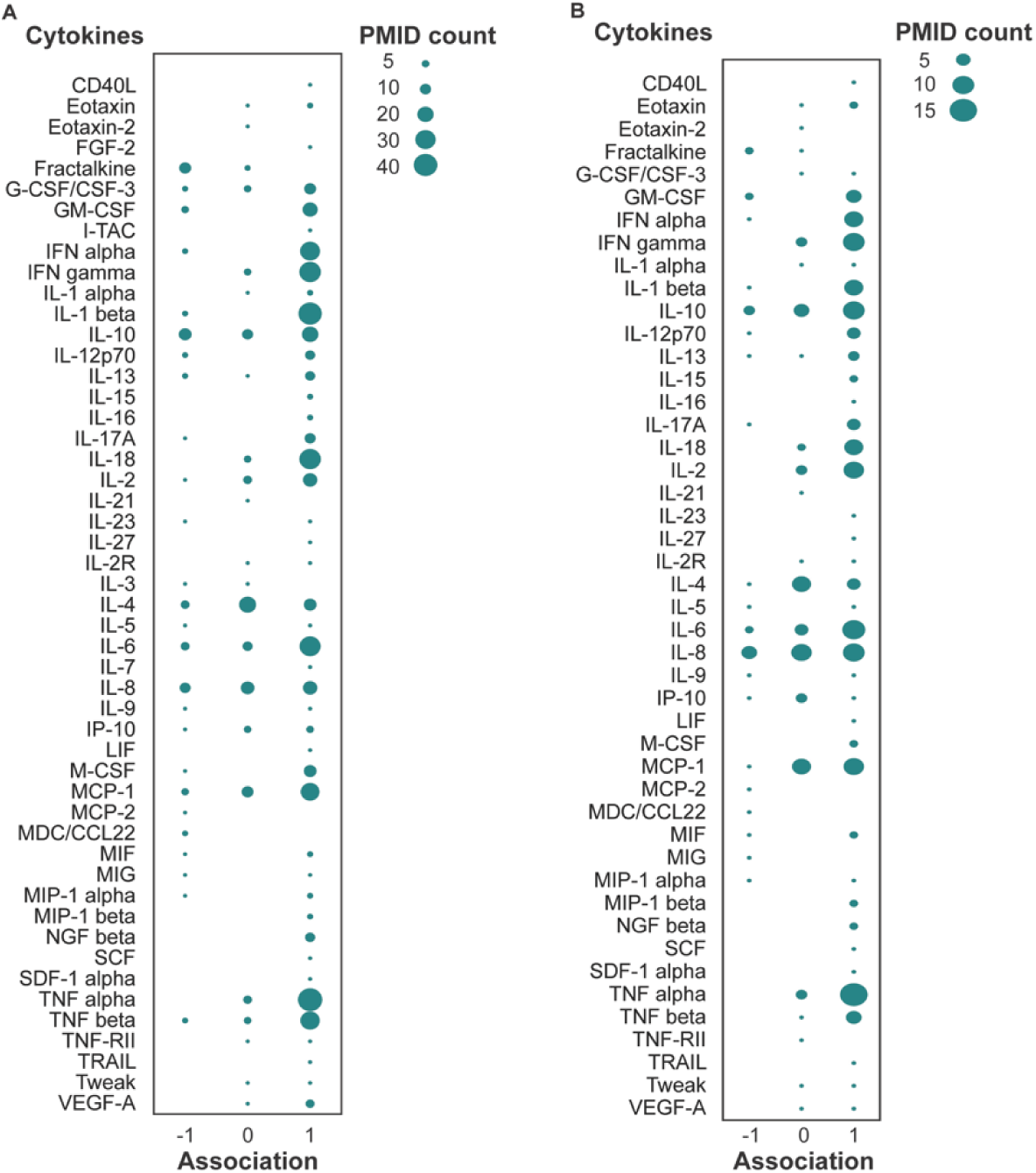
GPT-extracted primary literature overview of cytokines and PD. Scores of 1 were given to positive associations between higher cytokine concentrations and PD, −1 to negative associations between lower cytokine concentrations and PD, and 0 to no associations. Bubble size represents the number of papers supporting the association scores. A: Studies across various hosts, such as humans, animals, and cell lines B: Human studies only. Reviews, editorials, letters, and preprints were excluded, totaling 323 papers (out of 401) that were defined as primary literature used in this figure.

